# Chemical Rescue Serves as a Predictive Proxy for Glycosynthase Activity on Glycosidic Bonds via a Shared Glycosyl Oxocarbenium Transition State

**DOI:** 10.64898/2026.07.30.741823

**Authors:** Mohit Kumar, Chandra Kanth Bandi, Sri Vidya Vyjayanthi Tallavajhula, T. Emme Burgin, Srinivas V. S. Chakravartula, Shishir P. S. Chundawat

## Abstract

Engineered glycosynthases (GSs) are powerful biocatalysts for custom glycan synthesis, yet their optimization via directed evolution is severely constrained by bottlenecks in high-throughput screening for activated azido-sugar donors. Here, we demonstrate that chemical rescue (CR)—the azide-mediated restoration of hydrolytic activity in nucleophile-deficient mutants—serves as a predictive, high-throughput proxy for glycosynthase activity. Applying an azide-responsive *Escherichia coli* biosensor screen to a site-saturation mutagenesis library of *Thermotoga maritima α*-L-fucosidase (*TmAfc*), we established a strong rank-order correlation between CR and GS activities in both crude lysates (*ρ* = 0.73) and purified enzymes (*ρ* = 0.95). Transition path sampling and QM/MM umbrella sampling revealed that both pathways proceed through a shared oxocarbenium-ion-like transition state (Δ*G*^‡^ ≈ 8.7 kcal/mol), providing a structural and thermodynamic rationale for using CR to select for transition-state-stabilizing mutations. Biochemical characterization of top-performing variants yielded an engineered fucosynthase (TmAfc_D224G_N70D_T392S) exhibiting a nearly 100-fold enhancement in *V*_max_ alongside altered regioselectivity. This two-tiered screening framework leverages cost-effective chemical rescue assays to streamline glycosynthase engineering for tailored glycans synthesis.

## Introduction

Glycans are ubiquitous biopolymers that are essential for a broad range of biological processes^1,2^. They regulate cell-cell recognition, immune modulation, host-pathogen interactions, and cancer progression^3–8^. The precise three-dimensional structure of glycans can dictate its function, enabling their usage in biomedical research, diagnostics, and glycan-based therapeutics^8,9^.

Despite their critical importance for biomedical research and other applications, glycan synthesis remains chemically challenging^10^. Traditional chemical synthesis suffers from low yields, laborious multi-step procedures, and difficulties to achieve stringent regio- and stereoselectivity, which are required for biological activity^10,11^. Moreover, extensive protecting group strategies can limit scalability^12,13^

Enzymatic synthesis presents an alternative strategy to address some of the challenges^14^. Glycosyltransferases (GTs), which mediate glycan biosynthesis, offer high specificity for assembling complex oligosaccharides^15,16^. However, their applicability is constrained by several factors: many GTs are membrane-associated^16–18^ and have poor stability, exhibit limited substrate scope, and require costly, unstable nucleotide-sugar donors as substrates^19^. These challenges pose major economic barriers for the widespread use of GTs in industrial applications^15,18,20,21^.

An alternative enzymatic strategy focuses on repurposing Glycoside Hydrolases (GHs). Under certain reaction conditions, some GHs can perform *trans*-glycosylation reactions very efficiently^22,23^, and have been used for synthesis of complex glycans like Human Milk Oligosaccharides (HMOs)^24^. The structural diversity and ease of recombinant expression make GHs attractive for tailored engineering^25,26^. However, the dominant catalytic activity of wild-type GHs is hydrolysis of glycosidic bonds, and the yields of *trans*-glycosylation reaction products are typically low^27^.

This limitation motivated the development of "glycosynthases" (GSs)^27^, created by mutating the catalytic nucleophile in mutant GHs to suppress hydrolysis activity while enabling glycosyl transfer using activated sugar donors like glycosyl azides and fluorides ^28,29^. Although GSs offer a promising alternative to GTs, optimization of their performance is often a major bottleneck. Most efforts to optimize GSs are limited by bottlenecks in high-throughput screening since GS reactions typically lack simple optical readouts ^30^ and most assays for some donors (e.g., glycosyl azides or azido-sugars) are inherently low throughput^31^. As a result, rational design of GSs remains still underdeveloped for azido sugar targeting enzymes, while directed evolution efforts based on rational-selection followed by high-throughput screening is difficult for screening large mutant GS libraries^13^.

Given the limitations of existing GS assays, chemical rescue (CR) assays provide an alternative proxy for the biochemical characterization of GS mutants. During CR, exogenous nucleophiles can partially restore the hydrolytic activity of nucleophile-deficient mutant enzymes, resulting in reaction rates that depend on substrate binding and transition-state stabilization^32,33^. On the contrary, GS assays directly measure glycosidic bond formation between an activated donor and an acceptor sugar^34^. Although both phenomena depend on the same catalytic machinery, a systematic demonstration of a direct relationship between CR and GS, together with a mechanistic rationale for such a relationship, is still lacking.

Here, we employed a high-throughput azide biosensor platform for quantitative measurement of GS activity, and we applied it to a site-saturation library of a model GH29 α-L-fucosidase to enable a large-scale comparison of CR and GS activities. Our results suggest a positive correlation between CR and GS activities, prompting mechanistic analysis of both reaction pathways using quantum mechanics/molecular modeling (QM/MM) simulations, which indicated that both reactions proceed through closely related glycosyl oxocarbenium-ion-like transition states. Together, these findings suggest that CR activity can be used as a predictor of GS activity and provide a mechanistic foundation for cost-effective, high-throughput engineering of glycosynthases.

## Results & Discussion

### High-Throughput Biosensor Platform Design for Screening Fucosynthase Activity

In this study, we used the *TmAfc* D224G fucosynthase as a model enzyme to screen both glycosynthase and chemical rescue activity and develop a quantitative understanding of the relationship between the two reaction mechanisms (**Figure 1**). To evaluate the catalytic activities of *TmAfc* D224G fucosynthase variants spanning a broad range of reaction rates, a site-saturation mutagenesis (SSM) library was constructed targeting residues located within a 10 Å radius of the primary D224G mutation. In addition, a few variants previously reported by our group were also included in the library^13^. The resulting variants were then expressed in our *E. coli* biosensor strain and screened in a 96-well plate format to screen for two distinct activities for each variant. CR activity was measured by monitoring the release of p-nitrophenol (pNP) from the substrate pNP-Fucose in the presence of 2M sodium azide as exogenous nucleophile. GS activity was measured using the azide-responsive biosensor (see below for details), with β-fucosyl azide serving as the donor and pNP-Xylose as the acceptor.

**Figure 1.**
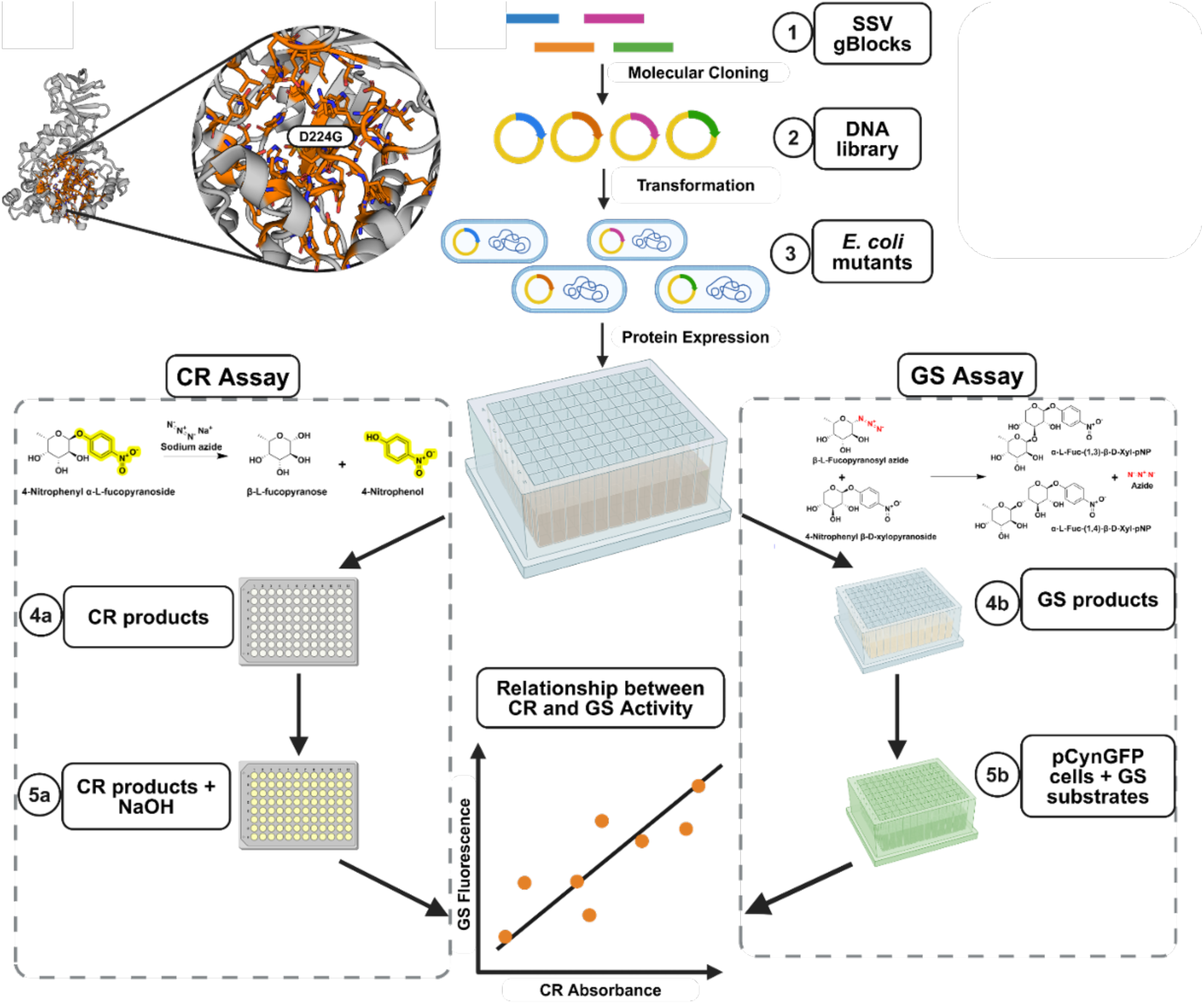
Schematic overview of the site-directed saturation mutagenesis workflow and high-throughput assays for monitoring chemical rescue (CR) and glycosynthase (GS) activities. The *TmAfc* structure (PDB ID: 2ZXD) shows residues within 10 Å of D224G targeted for mutagenesis. Resulting variants were expressed in E. coli and screened using CR and GS assays. In the CR assay, reaction between pNP-Fucose and azide produced p-nitrophenol, which was quantified by absorbance at 410 nm. In the GS assay, reaction between β-fucosyl azide and pNP-Xylose released azide, which activated a pCyn_v2_GFP biosensor, enabling GS quantification via GFP fluorescence (Ex 488 nm; Em 525 nm). The workflow revealed a positive relationship between CR and GS activities.

Previously, our group engineered an *Escherichia coli* strain harboring the pCyn_v2_GFP plasmid, which encodes a modified cyn operon (originally targeting cyanate) that exhibits high-sensitivity towards structurally analogous azide ion (N^3-^). This biosensor converts intracellular azide accumulation into a quantifiable fluorescence signal, as previously reported^31^. The biosensor design is well aligned with the catalytic mechanism of the *TmAfc* D224G fucosynthase. Briefly, during fucosyl transfer, the enzyme uses β-L-fucopyranosyl azide as the glycosyl donor and transfers the fucosyl moiety to an acceptor sugar group, such as p-nitrophenyl-β-D-xylose (pNP-Xylose), releasing one molecule of azide ion for each molecule of product formed. Because azide release is mechanistically coupled and stoichiometrically linked to glycosidic bond formation, GFP fluorescence provides a direct, real-time, and quantitative readout of fucosynthase activity within the cellular environment. Importantly our strategy circumvents many of the challenges associated with other screening formats, such as the need for product-specific sensors or artifacts arising from substrate or product by other cellular pathways. The robustness of this assay makes it a powerful and broadly applicable platform for screening glycosynthase systems that utilize an azide-based sugar donor.

### Whole-Cell Lysate Screen of Fucosynthase Variants Shows Strong Correlation Between Chemical Rescue and Glycosynthase Activity

Screening terminal catalytic activity of fucosynthases present in whole cell lysates of the generated library showed a positive correlation between normalized CR and GS values across the panel of enzyme mutants (**Figure 2a**). This result confirmed that enzyme variants that exhibited high CR activity consistently display high GS activity (Spearman correlation coefficient = 0.73). We interpret the non-linearity of the correlation with GS activity plateauing at the highest levels of CR activity as mechanistically informative. While the CR assay reflects the intrinsic catalytic capacity of the enzyme to stabilize the transition state, the whole-cell GS assay is a slightly more complex system. It is plausible that for the most active enzyme variants, overall reaction rates are limited by other biological processes, such as inter-membrane substrate transport or the saturation of the biosensor’s dynamic range. This would partly explain the observed leveling-off effect and enzyme performance within a cellular context. Altogether, the overall correlation between CR and GS activities provides strong evidence that these two reaction activities are mechanistically linked and supports the use of CR activity as a predictive metric in a screening framework.

**Figure 2.**
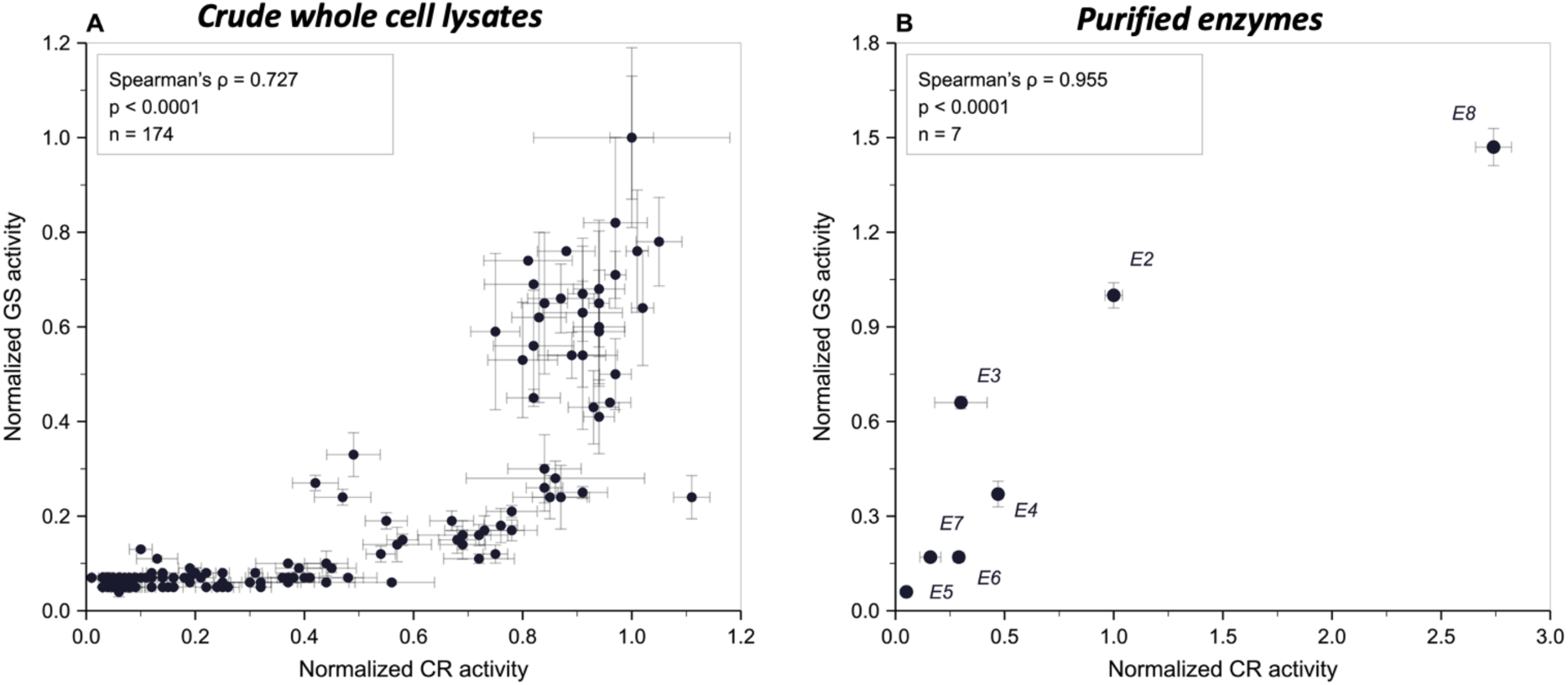
Correlation between Chemical Rescue (CR) and Glycosynthase (GS) activities in (A) crude whole cell lysates and (B) purified enzymes. (A) Relationship between normalized CR and GS activities for the site-saturation mutagenesis library measured in crude cell lysates using pNP-Xylose as the acceptor. Normalized CR values are plotted against normalized GS values, and the strength of association is quantified by a Spearman correlation coefficient of ρ = 0.727. Error bars represent the standard error of the mean for both measurements. (B) Relationship between normalized CR and GS activities quantified using HPLC for purified enzyme variants. The association between CR and GS values is summarized by a Spearman correlation coefficient of ρ = 0.955 (n = 7; the wild-type enzyme E1, which shows no glycosynthase activity, is not included). Error bars for CR and GS activity represent the standard deviations across measurements.

### Purified Fucosynthase *In-vitro* Assays Validated CR vs. GS Activity Correlation

To validate the findings from the cell lysate screen and biochemically characterize the engineered enzymes, we selected eight enzymes/mutants (E1-E8) spanning a broad range of activity levels. Variants E5 and E7 exhibited low CR and GS signals in the biosensor screen, whereas E3 and E4 displayed intermediate activities, in contrast, E8— a previously characterized high-activity variant included here as a high-end activity benchmark — displayed high CR and GS activity (protein sequences are listed in the supplementary document). These variants, together with the wild type glycosidase (TmAfc_Wt, E1), derived parent fucosynthase (TmAfc_D224G, E2), and additional promising candidates from the library, were expressed and purified to homogeneity by immobilized metal affinity chromatography (IMAC) (**Supplementary Figure S1**). Purified enzymes were then systematically assayed *in vitro* for both CR and GS activity. The strong correlation observed in the crude cell lysate screen was further verified with the purified proteins (Spearman correlation coefficient = 0.95) (**Figure 2b**), further supporting our preliminary *in vivo* screening assay results.

### Biochemical Characterization Reveals Catalytic Trade-Offs for Chemical Rescue

We performed detailed kinetic analyses of the CR reaction for purified enzyme variants exhibiting the highest CR activity by measuring the initial rates of pNP release at varying concentrations of the pNP-Fucose substrate (**Figure 3**; **Supplementary Figure S2**). The resulting data was fitted with the Michaelis-Menten model equation to extract kinetic parameters for the enzymatic reactions (**Table 1**; **Supplementary Figure S3**). This analysis revealed a striking result for mutant E8 (TmAfc_D224G_N70D_T392S). Compared to the parent E2 (TmAfc_D224G) glycosynthase, E8 exhibited a nearly 100-fold increase in V_max_, indicating a dramatic enhancement in its maximum catalytic rate. Interestingly, this was also accompanied by an increase in its K_m_ value, suggesting a slightly weaker affinity for the substrate. A combination of weaker ground-state binding and higher turnover has been reported for other improved glycosynthases, in which product release rather than substrate binding becomes rate-limiting^35^. These observations suggest that the improvements are not simply due to tighter substrate binding but rather arise from mutations that enhance the underlying chemistry of the catalytic reaction. This enhancement may stem from improved stabilization of the high-energy transition state, resulting in a substantial increase in catalytic turnover number (k_cat_), and consequently V_max._

**Figure 3.**
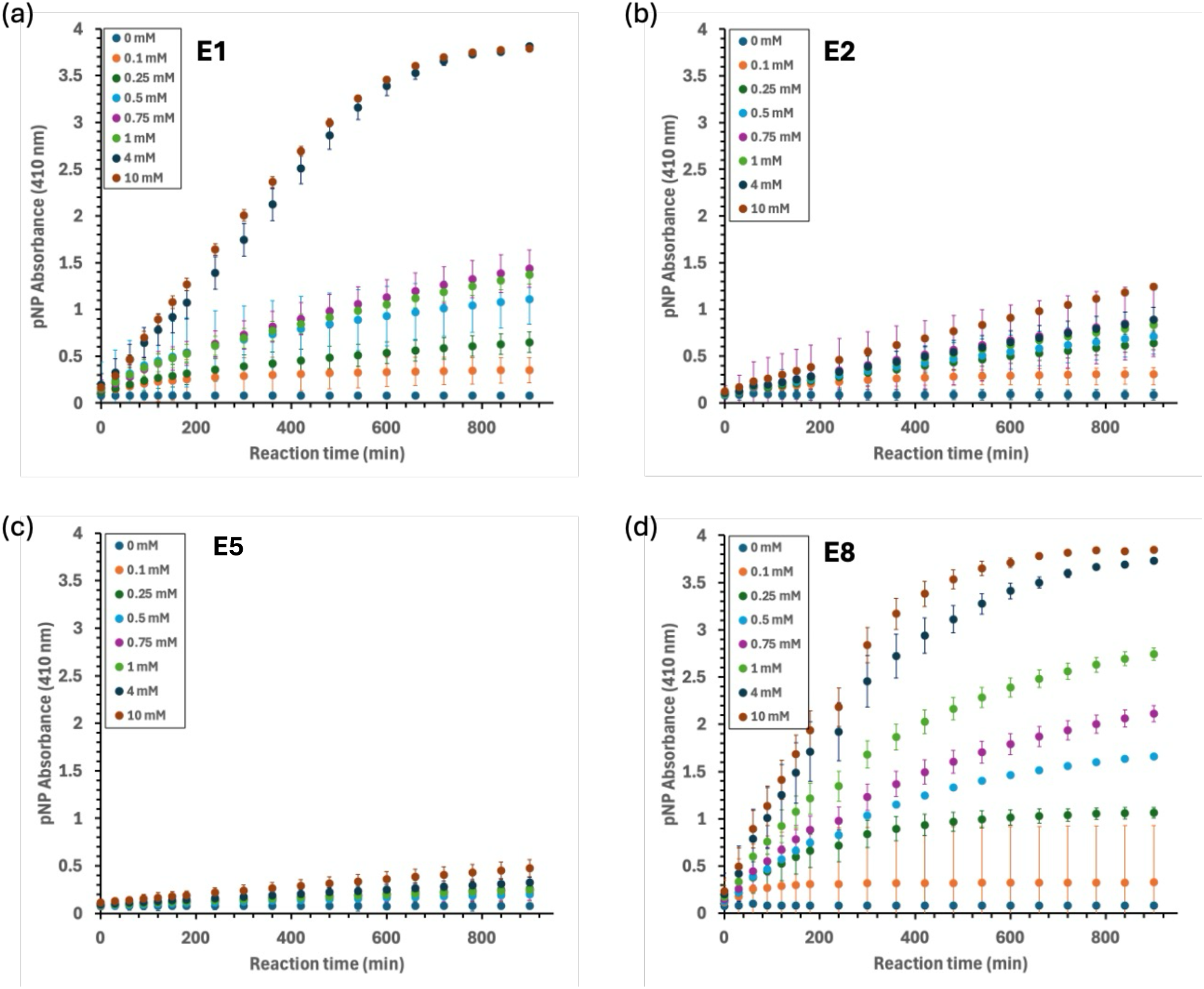
Kinetics of chemical rescue (CR) activity for purified enzyme variants. Reaction progress curves showing time-dependent release of p-nitrophenol (pNP), monitored at 410 nm, across pNP-Fucose concentrations ranging from 0–10 mM. Error bars represent standard deviations from replicate measurements. The different panels correspond to different enzyme variants E1 (a), E2 (b), E5 (c), and E8 (d). Panel (a) corresponds to the wild-type enzyme (E1), whose activity reflects native hydrolysis rather than azide-dependent chemical rescue and is shown for reference.

**Table 1:** Kinetic parameters of select *TmAfc* fucosynthase variants for Chemical Rescue (CR) reaction. Kinetic parameters were determined by fitting initial reaction rate data from the chemical rescue assay with varying pNP-Fucose concentrations to the Michaelis-Menten equation. Errors represent the standard deviation of the fitted parameters. Parameters for variant E5 derive from a lower confidence fit (R² = 0.57) and are reported for completeness only.

| Mutants | Vmax [M/s] | Km [M] | R<sup>2</sup> values |
| --- | --- | --- | --- |
| E1 | $2.67 \times 10^{-8} \pm 0.20 \times 10^{-8}$ | $2.76 \times 10^{-3} \pm 0.51 \times 10^{-3}$ | 0.9848 |
| E2 | $2.89 \times 10^{-10} \pm 0.32 \times 10^{-10}$ | $2.69 \times 10^{-4} \pm 1.22 \times 10^{-4}$ | 0.8545 |
| E5 | $9.83 \times 10^{-10} \pm 1.85 \times 10^{-10}$ | $2.11 \times 10^{-4} \pm 1.77 \times 10^{-4}$ | 0.5693 |
| E8 | $2.87 \times 10^{-8} \pm 0.10 \times 10^{-8}$ | $1.15 \times 10^{-3} \pm 0.13 \times 10^{-3}$ | 0.9916 |

### Top-Performing Fucosynthases Can Synthesize Fucosylated Disaccharides

To verify that the engineered enzymes identified from the CR/GS reactions screening catalyze the intended glycosylation reaction, glycosynthase (GS) reaction mixtures were analyzed by HPLC followed by ESI-MS analysis. Reactions containing the parent fucosynthase (*TmAfc_D224G*) E2 (positive control) and the mutant variants (E3-E8) produced two product peaks that were absent in the reactions with the wild-type enzyme (negative control), which retains the catalytic nucleophile and fails to catalyze GS reaction (**Supplementary Figure S4**). The highly active E8 variant also produced higher amounts of GS products compared to E2 (**Supplementary Table 1**). Mass spectrometric analysis of these peaks confirmed that they correspond to fucosylated disaccharides, specifically the α-L-Fuc-(1,4)-β-D-Xyl-pNP and α-L-Fuc-(1,3)-β-D-Xyl-pNP regioisomers (**Supplementary Figure S5, S6 S7 and Table 2**).

The relative proportion of the two regioisomers also varied across variants, shifting from a modest preference for the α-(1,4) product in the parent E2 toward the α-(1,3) product in E8, indicating that the selected mutations influence regioselectivity in addition to overall catalytic activity **(Supplementary Figure S5)**. Although the formation of multiple regioisomers reflects a common challenge in glycosidase-catalyzed synthesis reactions that should be addressed in future work, the analytical data overall confirms that our screening platform allows the engineering of efficient glycosynthase enzymes.

### A Shared Oxocarbenium-like Transition State That Mediates CR and GS

ESI-MS analysis of the chemical rescue reaction mixtures **(Supplementary Figure S8)**, performed alongside standards **(Supplementary Figure S9)**, confirmed the formation of β-fucosyl azide, an activated donor capable of supporting GS activity. This observation, together with the strong positive correlation between CR and GS activities, led us to hypothesize that both reactions proceed through a shared oxocarbenium-ion-like transition state, as previously proposed for GS reaction^36^ but not examined in the context of CR reaction. In this model, CR can be viewed as the reverse of GS: the GS reaction uses a hydroxyl group from the acceptor sugar, pNP-xylose in our system, as the nucleophile, whereas CR uses an external azide. Thus, mutations that stabilize the shared high-energy transition-state-like intermediate would be expected to enhance both reactions, consistent with our experimental results.

To test this hypothesis, we modelled the CR reaction mechanism of the D224G mutant using transition path sampling (TPS). TPS enables unbiased identification of reactive trajectories without assuming a reaction coordinate (RC) in advance, allowing statistically rigorous sampling of pathways that connect reactants and products (see Methods section for a detailed description of the methodology). Consistent with our hypothesis, the TPS results revealed that the CR reaction proceeds through an oxocarbenium-like transition state, closely resembling the GS mechanism (**Figure 4a-j**). We optimized the RC using inertial likelihood maximization^37^, drawing from a broad library of geometric features or collective variables (CVs) in the enzyme active site. These include distances between atoms among which chemical bonds form or break in the products compared to the reactants. The final RC is defined based on 3 collective variables (CV) shown in **Figure 5**: CVs 1 and 2 capture differences in distances between atoms where bonds form or break, and CV3 describes an angle. A detailed description of the methodology used to obtain the RC can be found in the Methods section. The optimized relationship between the RC and the 3 CVs was RC = -6.054 – 2.663*CV1 – 0.218*CV2 + 0.031*CV3.

**Figure 4.**
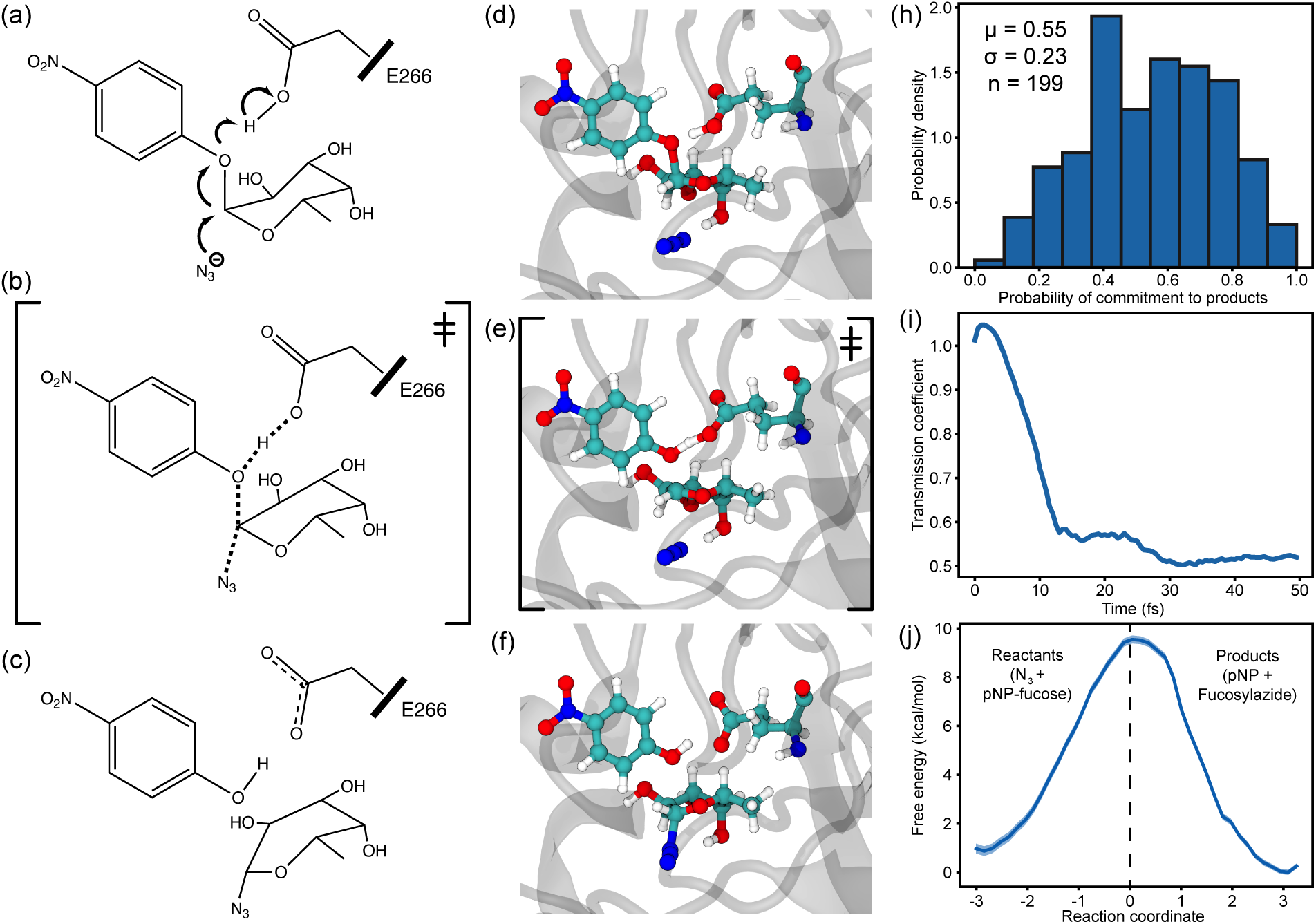
Transition path sampling (TPS) characterization of the chemical rescue mechanism. (a–c) Schematic snapshots of the proposed CR reaction pathway: reactant state with azide approaching pNP-fucose, an oxocarbenium-like transition state with partial bond formation and proton transfer involving Glu266, and the product state with azide bound at the anomeric carbon. (d–f) All-atom representations of a representative reactive trajectory corresponding to panels a–c. (h) Committor analysis results showing a near-centered distribution, supporting identification of a valid transition-state ensemble. The number of independent sets of initial coordinates is n, µ is the distribution mean, and σ is the standard deviation. (i) Transmission coefficient from committor simulations, plateauing at ∼0.52. (j) Free-energy profile along the optimized reaction coordinate obtained from umbrella sampling. Shaded region represents standard error of the mean. Transition state is indicated by a dashed line.

**Figure 5.**
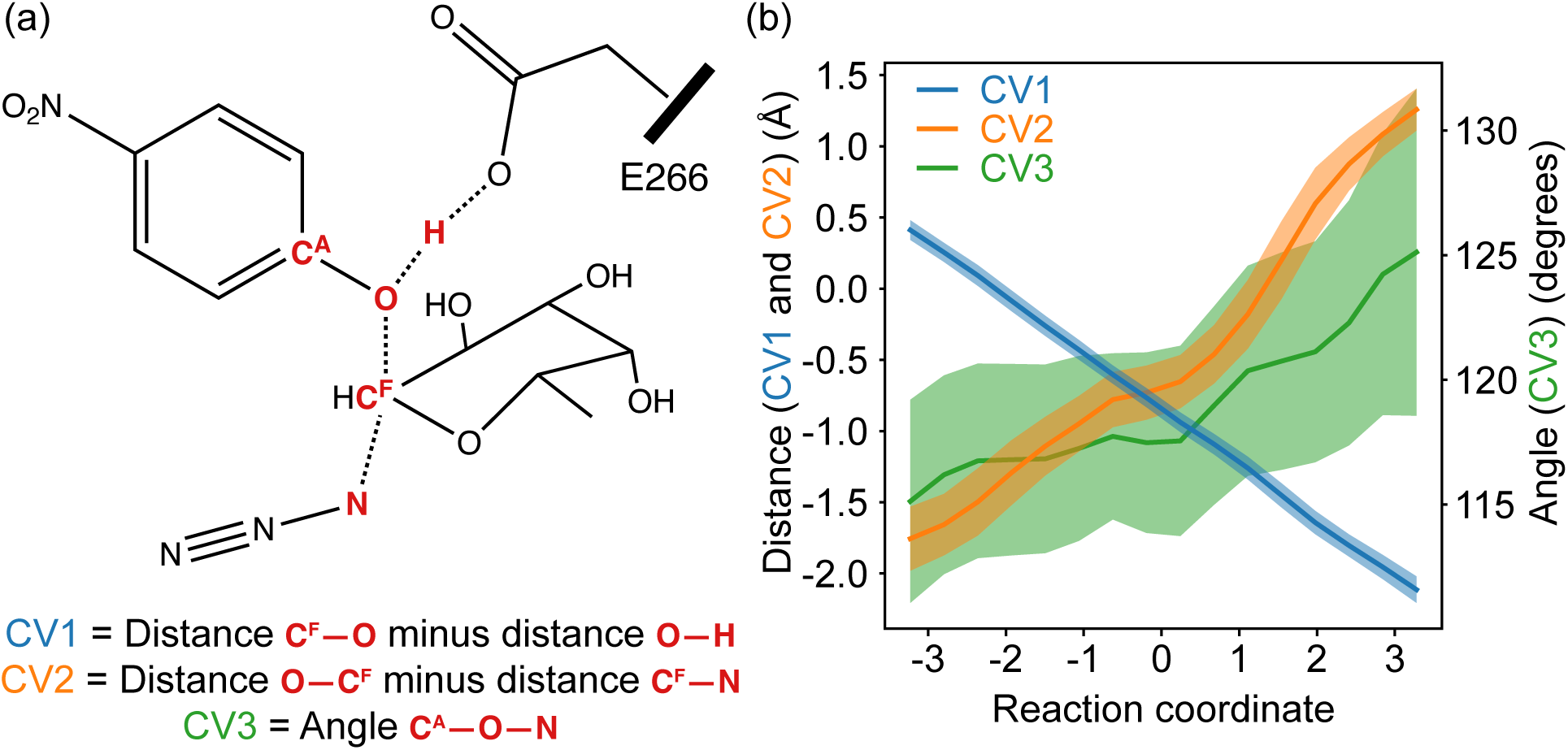
Collective variable (CV) definitions and behavior along the reaction pathway. (a) Diagram of the CVs used in the optimized reaction coordinate. Atoms contributing to each CV are shown in red. CV1 and CV2 track differences in key bond-forming and bond-breaking distances, while CV3 corresponds to an angle formed by a partially discontiguous set of atoms. (b) Average CV values across reactive trajectories obtained from transition path sampling. Shaded regions indicate one standard deviation.

To validate this proposed RC, we performed committor analysis starting from nearly 200 independent configurations near the predicted transition state, as described in Methods (**Figure 4h**). These simulations showed an approximately 50% likelihood of progressing to the products state, indicating that the coordinate accurately describes transition-state configurations. Umbrella sampling along this RC produced a smooth free-energy profile (ΔG^‡^ ≈ 8.7 kcal/mol; K_eq_ ≈ 4) (**Figure 4j**; **Supplementary Figure S11**). The resulting transmission coefficient along the RC was ∼0.52 (**Figure 4i**), and based on the Eyring equation, the estimated initial forward rate constant for this reaction is 7.1 × 10^6^ s⁻¹. Assuming a standard ±2 kcal mol⁻¹ uncertainty in the MD simulations puts the predicted range of initial rate constants ranges between 3.5×10^5^ − 1.5×10^8^ s⁻¹, with equilibrium constants between 0.12 − 50.

The computed rate constant describes the elementary chemical step of glycosyl–azide bond formation. That it is not directly comparable to the steady-state turnover measured over the assay time course, which additionally reflects substrate binding and product release, suggests that the chemical reaction itself may not be the rate-limiting step in the turnover of CR under the conditions tested. Altogether, the computed energetics support our hypothesis that CR and GS proceed through a common oxocarbenium-like transition state. This provides a mechanistic explanation for the positive correlation between CR and GS activities that we observed experimentally for our fucosynthase mutant library.

## Conclusion

In this work, we established a streamlined and cost-effective strategy for engineering glycosynthases, based on a strong rank-order correlation between their CR and GS reaction activities. This allows application of inexpensive CR assays to serve as a reliable high-throughput primary screen, substantially reducing reliance on specialized GS assays that require azido-sugar donors and complex analytical workflows thus directly addressing one of the key bottlenecks in GS engineering^38,39^. In our platform, crude cell lysates CR activity screen serves as a high-throughput filter that rapidly identifies promising variants with enhanced GS activity, while GS assays are reserved for the final biochemical and analytical validation of a smaller set of top candidates. This strategy lowers material costs, improves assay scalability, and facilitates glycosynthase engineering for basic research and engineering applications^13,40^.

Beyond its methodological advantages, this study provides key molecular insights into the CR catalytic reaction mechanism for the first time. The positive correlation between CR and GS activities observed experimentally provided initial evidence that both reactions proceed through a common transition state. Guided by previous mechanistic studies of the GS reaction^35,41^, we hypothesized that this common transition state is an oxocarbenium ion. Our computational analysis supports this hypothesis: transition path sampling and RC optimization revealed that the CR reaction follows the reverse pathway to that previously established for the GS reaction. Since this analysis was performed only for the parent D224G enzyme, for the engineered variants (e.g., E8), the shared-transition-state interpretation is inferred from the agreement between the computed CR pathway and the previously reported GS pathway, along with the experimental correlation, rather than from variant-specific simulations. On this basis, mutations that stabilize the oxocarbenium-like transition state would be expected to accelerate the turnover of both reactions.

Variant E8 exemplifies this principle on how increased fucosynthase GS activity can be explained by the stabilization of the CR transition state. Its elevated CR activity indicated stabilization of the oxocarbenium-like transition state, which translates into increased GS efficiency despite a modest reduction in ground-state substrate affinity (higher Km). These findings refine our understanding of the catalytic mechanism of glycosynthases and reinforces the idea that the CR reaction can serve not merely as a proxy, but also as a predictive probe of transition-state energetics during glycosidase engineering. Improving CR effectively selects for variants with enhanced transition-state stabilization, which translates into a more efficient catalyst for both reactions. These insights can guide future engineering strategies by providing a conceptual foundation for rational design strategies targeting transition-state stabilization as the key determinant of GS activity.

Although our study focuses on GH29 enzymes and a specific set of azido-sugar donors, the conceptual framework may extend to other retaining glycosidases. By substituting the acceptor sugar, our platform can also be tailored to discover or optimize other glycosynthases for specific targets, including high-value structures such as HMOs^42,43^, host-pathogen recognition^44^, and therapeutically relevant glycans^45^. Nonetheless, differences in active-state architecture, acceptor binding modes, or leaving-group chemistry might influence the CR-GS relationship, and further studies across diverse enzyme classes are required to determine the breadth of applicability of our screening approach.

## Materials and Methods

### Bacterial Strains, Reagents, and Plasmids

*Escherichia coli* strain E. cloni® 10G (Lucigen, WI) was used for all cloning experiments, and *E. coli* BL21(DE3) was used for protein expression. Standard chemical reagents were purchased from VWR, Thermo Fisher Scientific, or Sigma-Aldrich unless otherwise stated. Restriction enzymes and Phusion DNA polymerase were sourced from New England Biolabs (NEB). The plasmid pEC-TmAfc, containing the E. coli-codon-optimized gene for *Thermotoga maritima* α-fucosidase (Tm0306), was available from previous work. Site-saturation variant (SSV) gene fragments (gBlocks) were synthesized by Twist Bioscience (CA). Oligonucleotide primers were obtained from Integrated DNA Technologies (NJ), and DNA sequencing was performed by Genewiz (Azenta, NJ). Carbohydrate substrates, including 4-nitrophenyl α-L-fucopyranoside (pNP-Fucose) and p-nitrophenyl-β-D-xylose (pNP-Xylose), were procured from Carbosynth. β-L-fucopyranosyl azide was obtained from Chemily Glycosciences.

### Engineering and Preparation of Library Constructs

The gene encoding TmAfc (Tm0306) from the hyperthermophile *Thermotoga maritima* was previously cloned into a pEC vector suitable for expression in *E. coli*. The catalytic nucleophile Asp224 was mutated to Gly using standard site-directed mutagenesis protocols to create the parent fucosynthase construct (D224G). For the generation of the site-saturation mutagenesis library, residues within a 10 Å radius of position 224 were targeted. SSV gBlocks containing randomized codons at these positions were synthesized and cloned into the DpnI-digested pEC-TmAfc D224G backbone using Gibson Assembly® Master Mix (NEB) following the manufacturer’s protocol. The assembly reaction was transformed into E. cloni 10G competent cells, which were then plated on Luria-Bertani (LB) agar containing 50 µg/mL kanamycin. The diversity and quality of the library were confirmed by sequencing a representative number of colonies.

### Protein Expression and Purification

For crude cell-based assays, individual colonies from the library plates were inoculated into 96-deep-well plates containing LB medium with kanamycin and cultured overnight at 37°C. The next day, cells were harvested by centrifugation, and the pellets were resuspended in water to create cell suspensions for activity assays. For purified protein assays, E. coli BL21(DE3) cells were transformed with selected mutant plasmids. A 50 mL starter culture was grown overnight and used to inoculate 1 L of LB medium supplemented with kanamycin. The culture was cultured at 37°C with shaking at 200 rpm until the optical density at 600 nm (OD600) reached 0.4–0.8. Protein expression was then induced with 0.5 mM IPTG, and the culture was incubated for an additional 20 hours at 25°C. Cell pellets were harvested by centrifugation at 4,000 x g for 20 min. The pellet was resuspended in lysis buffer (20 mM sodium phosphate, 500 mM NaCl, 20% glycerol, pH 7.4) containing lysozyme and protease inhibitors, and cells were lysed by sonication. The soluble fraction was clarified by centrifugation and applied to a Ni-NTA IMAC column. The proteins, which contained a His tag at their C terminus, were eluted with an imidazole gradient. The purified protein was buffer exchanged into a storage buffer (10 mM MES, pH 6.0) using an ultrafiltration device. Protein concentration was determined by measuring absorbance at 280 nm using a SpectraDrop micro-volume slide, and purity was assessed by SDS-PAGE using stain-free gels imaged with a Gel Doc EZ Imager (Bio-Rad).

### Activity Assays

Details for enzyme assays and product analysis are provided below.

#### Chemical Rescue (CR) Assay

For purified proteins, 2 µg of enzyme was incubated in a 50 mM MES buffer (pH 6.0) with 2 mM pNP-Fucose and 2 M sodium azide at 60°C for 2 hours. For crude cell assays, cell suspensions were used under similar conditions. The reaction was quenched by adding 100 µL of 0.1 M NaOH to 100 µL of the reaction mixture, and the release of p-nitrophenol (pNP) was quantified by measuring absorbance at 410 nm with a SpectraMax M5e microplate reader. A standard curve of pNP was used for quantification.

#### Glycosynthase (GS) Assay

For purified proteins, 40 µg of purified enzyme was incubated with 10 mM β-L-fucopyranosyl azide and 50 mM pNP-β-D-xylose in 50 mM MES buffer (pH 6.5) at 60°C for 24 hours. Reaction progress was monitored by product analysis as previously described ^29,33^. For crude cell biosensor screening, a two-plate assay was used. In the "reaction plate," library transformants were grown, harvested, and resuspended. Substrates (10 mM β-fucosyl azide and 50 mM pNP-Xylose) were added, and the plate was incubated at 60°C. In the "reporter plate," *E. coli* cells containing the pCyn_v2_GFP biosensor plasmid were grown. After the GS reaction, an aliquot from the reaction plate was transferred to the reporter plate. After further incubation to allow for GFP expression, fluorescence was measured (excitation: 488 nm; emission: 525 nm) to quantify the amount of azide released, which is proportional to GS activity.

#### Kinetic Analysis

For kinetic parameter determination, the CR assay was performed with purified enzymes using a range of pNP-Fucose concentrations (0–10 mM). Initial reaction rates were determined by monitoring pNP absorbance at 410 nm over time. The data were fitted to the Michaelis-Menten equation using non-linear regression analysis in GraphPad Prism to determine V_max_ and K_m_ values.

#### Product Analysis

High-Performance Liquid Chromatography (HPLC): Product separation was performed on an Agilent LC system using a SUPELCOSIL LC-NH2 HILIC column (25 cm × 4.6 mm, 5 µm). A gradient of acetonitrile and water was used at a flow rate of 1 mL/min. UV absorbance was monitored at 254 nm and 300 nm.

### Electrospray ionization-Mass spectrometry (ESI-MS)

MES buffer controls and GS reaction products in MES buffer were diluted in 50% acetonitrile/water and infused by a manual syringe pump (Fisher Scientific, USA) into a quadrupole-TWIMS-TOF hybrid mass spectrometer (Synapt G2-Si qTOF; Waters Corporation, Manchester, UK) in positive ionization mode. The flow rate was adjusted to 600 µL/hr. MS spectra (100 - 600 m/z, 1 min acquisition) were acquired using the following settings: capillary voltage, 3.0 kV; sampling cone voltage, 40 V; source offset 30°C; source temperature, 100 °C; desolvation temperature, 250°C: cone gas, 50 L/Hr; desolvation gas, 600 L/Hr.

### Liquid Chromatography-Mass Spectrometry (LC-MS)

Sugars, nitrophenyl glycosides and glycosynthase reaction products were analyzed on a Waters Acquity H-class UPLC system coupled to the hybrid Xevo-G2-XS-TOF high-resolution mass spectrometer (HRMS) (Waters Corporation, Manchester, England). Chromatography was performed on the Acquity UPLC BEH Amide 2.1mmx100mm 1.7µm HILIC column (Waters Corporation). The column temperature was set at 40°C and the samples were kept at 10 °C. A sample volume of 2 µL was injected into the LC column. The composition of the mobile phase was 80:20 acetonitrile:water containing 0.1% ammonium hydroxide as eluent A, and 30:70 acetonitrile:water containing 0.1% ammonium hydroxide as eluent B. A step-gradient of increasing eluent B was applied at a flow rate of 0.2 mL/min: 0 – 2.0 min, 10% B; 2 – 4.0 min, 20% B; 4.1 – 6.0 min, 50% B; 6.1 – 10.0 min, 70% B; 10.1 – 12.0 min, 10% B. The needle wash solution was 10% – 90% MQ water. ESI-MS analysis was performed in negative ionization under sensitivity mode (100 - 1000 m/z) using the following settings: capillary voltage, 3.0 kV; sampling cone voltage, 30 V; source offset, 60°C; source temperature, 125°C; desolvation temperature, 250°C; cone gas flow rate, 50 L/hr; desolvation gas flow rate; 600 L/hr. Data analysis was performed using the Mass Lynx 4.1 software.

### Molecular dynamics simulations and transition path sampling

Transition path sampling (TPS) was conducted using the ATESA software package ^46,47^. Aimless shooting proceeded until the information error in the inertial likelihood maximization procedure fell below the threshold value of 0.1. Inertial likelihood maximization used ATESA’s default two-line test procedure to identify an RC with a balance between quality of fit to the aimless shooting data and a small number of CVs in the RC. The reactant and product states were defined according to the following criteria:

Reactants: Distance from the E266 acidic oxygen to E266 acidic hydrogen less than 1.15 Å; distance from fucose O1 to E266 acidic hydrogen greater than 1.5 Å; distance from fucose O1 to fucose C1 less than 2.0 Å; and distance from reactive azide N to fucose C1 greater than 2.2 Å.

Products: Distance from the E266 acidic oxygen to E266 acidic hydrogen greater than 1.5 Å; distance from fucose O1 to E266 acidic hydrogen less than 1.1 Å; distance from fucose O1 to fucose C1 greater than 2.5 Å; and distance from reactive azide N to fucose C1 less than 2.0 Å.

Aimless shooting was divided into 12 independent threads, each starting from the same initial transition state guess obtained using the *find_ts* routine from ATESA. New shooting moves originated from accepted trajectories, based on a randomly selected frame of either the forward or backward trajectory (also chosen randomly) between 1 and 15 frames from the previous starting frame.

Molecular dynamics (MD) simulations were conducted using Amber 22 ^48^. All simulation parameters, including the selection of the QM region and the QM method (DFTB3) used, were the same as in our previous work on the GS reaction of the same enzyme^36,49^. These included an Andersen thermostat with a time constant of 1000 steps (much longer than the length of all simulations) and a 0.5 fs simulation timestep^50^.

Umbrella sampling simulations were restrained along the RC using Plumed^51^. The simulations were divided into evenly spaced windows 0.25 units apart along the RC, with a restraint weight of 50 kcal/mol. Five independent simulations were performed in each restraint window. Samples were decorrelated from their initial values according to ATESA’s implementation of pyMBAR^52^. Histograms of sampling showing good overlap between adjacent windows are shown in **Supplementary Figure S10**.

## Acknowledgements

This research was supported by the National Science Foundation (NSF) grant 1904890 from CHE division (CLP program), Rutgers School of Engineering (SOE), and Rutgers Chemistry and Chemical Biology (CCB) Program. We extend our gratitude to the current and former members of the Chundawat lab for their invaluable assistance in our experimental endeavors.

