## Supplementary Material for "Chemical Rescue Serves as a Predictive Proxy for Glycosynthase Activity on Glycosidic Bonds via a Shared Glycosyl Oxocarbenium Transition State"

**Supplementary Methods:**

### Gene Synthesis

Gene synthesis and cloning were conducted on a model GH family 29 fucosidase enzyme from the hyperthermophile *Thermotoga maritima*, known as *Tm-alpha-fucosidase* (*TmAfc*). The native (wild-type) gene Tm0306 (locus tag TM0306; UniProt Q9WYE2) encoding TmAfc, was codon-optimized for expression in *E. coli* and custom synthesized with AsiSI and BamH1 restriction sites by Genscript Biotech Corporation in the pUC57 vector. The Tm0306 gene was subsequently sub-cloned from the pUC57 vector into our customized pEC vector, which features a T5 promoter and a Kanamycin selection marker, using standard restriction cloning techniques.

The catalytic nucleophile of Tm0306 (D224) was mutated to alanine (D224A), serine (D224S), and glycine (D224G) using standard site-directed mutagenesis protocols. For this, 0.5 µM of forward and reverse primers for mutagenesis were mixed with 20 ng of plasmid DNA in a 10 µl reaction volume. The reaction utilized 1X Master Mix (Phusion DNA polymerase, 200 µM dNTPs, 1X Phusion HF buffer, 1.5 mM MgCl2) with 5% DMSO, and the volume was adjusted to 10 µl with nuclease-free PCR water. Amplification was confirmed via DNA gel electrophoresis before the PCR products were digested with 10 U of Dpn1 enzyme (New England Biolabs) at 37°C for 1 hour. The Dpn1-digested mixture was transformed into E. Cloni 10G competent cells (Lucigen) using the Zymo transformation kit and plated on LB agar plates with the appropriate selection marker (Kanamycin). Several random colonies were picked, plasmid DNA was extracted and verified by DNA sequencing (Genewiz/Azenta).

### SSV gBlocks PCR

SSV gBlocks were procured from Twist Bioscience in a lyophilized form. Because gBlocks have overhangs, PCR amplification for insert preparation was unnecessary. For vector PCR, 0.5 µM of forward and reverse primers were combined with 20 ng of Tm0306_WT plasmid DNA, along with 0.5 µM of each mutagenesis primer. The reaction mix comprised 1X Master Mix (Phusion DNA polymerase, 200 µM dNTPs, 1X Phusion HF buffer, and 1.5 mM MgCl2), supplemented with 5% DMSO, and adjusted to a final volume of 10 µl using nuclease-free PCR water. The standard PCR conditions used for amplification of the insert were used. Once PCR was complete, 2 µl of the PCR product was mixed with 3 µl PCR water and 1 µl of purple loading dye and run in a SYBR safe DNA gel alongside 5 µl of DNA ladder at 120 V for 45 minutes. The remaining PCR products were purified using a PCR extraction kit from IBI Scientific. Next, 2 µl of the purified PCR product was used in spectrophotometer to measure DNA concentration.

Gibson cloning, and transformation**:** Reaction mixtures were prepared for Dpn1 digestion along with 1X Cut smart buffer in a total reaction volume of 10 µl and digested with 20U of Dpn1 at 37°C for 1 hour. After Dpn1 digestion, 1.5U of Gibson cloning reagent was added along with NEB buffer 2.1 was added to the PCR reaction mixture in a total reaction volume of 20 µl and incubated at 37°C for 45 minutes. The PCR products along with appropriate inserts were then placed on ice immediately and transformed into E. cloni 10G cells and incubated at 37°C for 2 hours. The transformation mixture was plated on LB-agar plates with 50 µg/ml kanamycin and incubated at 37°C for 16 hours. Several colonies were observed on the LB agar plates, and colony screening was performed to identify the correct colonies.

Colony Screening**:** For colony screening, 20 random colonies were picked from the transformed plate. The tip used to pick each colony was also transferred to LB media with 50 µg/ml kanamycin and incubated at 37°C for 14-15 hours. The overnight grown culture was centrifuged next day, and plasmid extraction was done, and all 20 samples were sent for sanger sequencing using 0.5 µM NcoI forward (TTGCTTTGTGAGCGGATAAC) and 0.5 µM T7 terminator reverse (GCTAGTTATTGCTCAGCGG) primers. The analysis of all colony and PEDEL-AA analysis was done from this data.

**Protein expression and purification**

Sequence-verified wild-type (Tm0306_WT) and corresponding nucleophile mutant (Tm0306_D224A/S/G) along with other desired mutants' DNA plasmids were transformed into *E. coli* BL21 (DE3) competent cells and plated onto LB agar plates containing 50 µg/ml kanamycin. Individual colonies were selected for inoculation in a 50 ml starter culture of LB media supplemented with 50 µg/ml kanamycin and incubated at 37°C for 12-16 hours. The overnight cultures were then transferred into 1000 ml LB media with 50 µg/ml kanamycin and grown at 37°C until the culture density reached an OD600 of 0.4-0.8. Protein expression was induced with 0.5 mM Isopropyl β-D-1-thiogalactopyranoside (IPTG), and the cultures were incubated at 25°C for 20 hours. Cell pellets were collected by centrifugation and stored in the freezer until needed. These pellets were resuspended in lysis buffer (20 mM sodium phosphate, 500 mM NaCl, and 20% glycerol, pH 7.4) at a 1:5 ratio of cells to buffer solution (total weight basis), along with a protease inhibitor cocktail (1 µM E-64, 0.5 mM benzamidine, and 1 mM EDTA) and lysozyme (10 µg/ml), and lysed by sonication on ice. The lysed pellets were centrifuged, and the supernatant containing the soluble protein was collected. C-terminal his-tagged proteins were purified from other E. coli proteins using an IMAC (Ni-immobilized metal affinity chromatography) column with the NGC-FPLC system (Bio Rad). The Ni-IMAC column was first equilibrated with IMAC binding buffer (100 mM MOPS, 10 mM imidazole, 500 mM NaCl, pH 7.4). The cell lysate supernatant was then loaded onto the column, and IMAC binding buffer was run through to remove non-specifically bound proteins. The protein of interest was eluted with IMAC elution buffer (100 mM MOPS, 500 mM imidazole, 500 mM NaCl, pH 7.4). The eluted protein was buffer-exchanged using desalting columns (GE Healthcare, Catalog number: 17-0851-01) into 10 mM of 2-morpholin-4-ylethanesulfonic acid (MES) at pH 6. The concentration of purified protein was estimated using the Spectradrop UV spectrophotometer (SpectraMax M5e) based on 280 nm absorbance. The purity of all enzymes was confirmed by SDS-PAGE (see Supplementary Figure) through gel densitometric analysis using pre-cast stain-free (Bio-Rad) protein electrophoresis gels.

Protein sequences for E1 and E8 mutant (taken from previous work in our lab) is given here, and E2, E3, E4, E5 and E7 were also from wild type as following: E2 (TmAfc_D224G), E3 (TmAfc_D224G_V53Y), E4 (TmAfc_D224G_L50V), E5 (TmAfc_D224G_M225R), E6 (TmAfc_D224G_N270H), E7 (TmAfc_D224G_L191M).

1) E1 (TmAfc_Wt)

MISMKPRYKPDWESLREHTVPKWFDKAKFGIFIHWGIYSVPGWATPTGELGKVPMDA
WFFQNPYAEWYENSLRIKESPTWEYHVKTYGENFEYEKFADLFTAEKWDPQEWADLF
KKAGAKYVIPTTKHHDGFCLWGTKYTDFNSVKRGPKRDLVGDLAKAVREAGLRFGVYY
SGGLDWRFTTEPIRYPEDLSYIRPNTYEYADYAYKQVMELVDLYLPDVLWNDMGWPEK
GKEDLKYLFAYYYNKHPEGSVNDRWGVPHWDFKTAEYHVNYPGDLPGYKWEFTRGI
GLSFGYNRNEGPEHMLSVEQLVYTLVDVVSKGGNLLLNVGPKGDGTIPDLQKERLLGL
GEWLRKYGDAIYGTSVWERCCAKTEDGTEIRFTRKCNRIFVIFLGIPTGEKIVIEDLNLS
AGTVRHFLTGERLSFKNVGKNLEITVPKKLLETDSITLVLEAVEE

2) E8 (TmAfc-D224G-N70D-T392S)

MISMKPRYKPDWESLREHTVPKWFDKAKFGIFIHWGIYSVPGWATPTGELGKVPMDA
WFFQNPYAEWYEDSLRIKESPTWEYHVKTYGENFEYEKFADLFTAEKWDPQEWADLF
KKAGAKYVIPTTKHHDGFCLWGTKYTDFNSVKRGPKRDLVGDLAKAVREAGLRFGVYY
SGGLDWRFTTEPIRYPEDLSYIRPNTYEYADYAYKQVMELVDLYLPDVLWNGMGWPEK
GKEDLKYLFAYYYNKHPEGSVNDRWGVPHWDFKTAEYHVNYPGDLPGYKWEFTRGI
GLSFGYNRNEGPEHMLSVEQLVYTLVDVVSKGGNLLLNVGPKGDGTIPDLQKERLLGL
GEWLRKYGDAIYGTSVWERCCAKTEDGTEIRFTRKCNRIFVIFLGIPSGEKIVIEDLNLS
AGTVRHFLTGERLSFKNVGKNLEITVPKKLLETDSITLVLEAVEE

**Protein characterization**

SDS-PAGE was performed for all purified proteins used for validation and characterization and marked against the 54 KDa spot from the ladder as shown in Supplementary Figure S1. All proteins were quantified using spectrophotometry (SpectraMax). Additionally, band intensity percentage was calculated for proportional quantification.

**Fucosidase activity and chemical rescue assays**

The activity of the purified enzymes E1 to E8 were evaluated using pNP-F (4-nitrophenol α-fucopyranoside) as the substrate, obtained from Carbosynth Limited. Briefly, 2 μg of protein was added to 2 mM pNP-F in a reaction buffer containing 50 mM MES at pH 6 and incubated at 60°C for 1.5 hours. Blank wells with only pNP-F and buffer, without added proteins, served as controls. Each reaction mixture was prepared in triplicate. After 1.5 hours, 100 μl of the reaction mixture was transferred to a transparent 96-well microplate along with 100 μl of 1 M NaOH, and the absorbance was measured at 410 nm using a UV/Vis spectrophotometer (SpectraMax M5e) to determine the total released pNP absorbance from substrate hydrolysis. A pNP calibration curve was constructed to correlate measured absorbance with estimated concentration. To recover or 'rescue' the hydrolytic activity of the inactive nucleophile mutants, high concentrations of external nucleophiles like sodium azide and sodium formate (2 M each) were added to reaction mixtures and incubated at 60°C for 2 hours. After the reaction was completed, 30 µl of the reaction mixture was transferred to a transparent 96-well microplate, mixed with 70 µl of DI water and 100 µl of 0.1 M NaOH, and the absorbance was measured at 410 nm using a UV/Vis spectrophotometer (SpectraMax M5e).

**HPLC analysis of GS reaction products**

Hydrophilic interaction (HILIC) chromatography was performed using Agilent Online LC system (G3167A, Agilent Technologies Inc., Germany), that consisted of a flexible pump, online sampler manager, a thermostated column compartment and a diode array detector. The HPLC was performed in a gradient mode. The mobile phase consisted of acetonitrile and water, which was employed at the flow rate of 1 mL/min. Briefly, the gradient consisted of 100% acetonitrile for 0-5 minutes, followed by 10% water for 5-10 minutes, 35% water for 10-25 minutes, and 100% acetonitrile for 25-30 minutes. The separation of mono and disaccharide products was analyzed using the SUPELCOSIL LC-NH2 HPLC column (120 Å, 25 cm × 4.6 mm, 5 µm; Supelco Inc., USA) with the column compartment at 60°C. The sample injection volume was 2 µL and the autosampler compartment was maintained at 40°C. UV absorbance peaks were detected at 254±10 nm & 300±10 nm with the reference wavelength at 360±100 nm. Then, 5 µl of the reaction mixture was injected onto the column, and all pNP-based products (i.e., pNP-xylose, α-L-Fuc-(1,4)-β-D-Xyl-pNP, and α-L-Fuc-(1,3)-β-D-Xyl-pNP) were detected using a DAD detector at 254 nm and 300 nm wavelengths. Raw data were acquired and analyzed using the Shimadzu LabSolutions software. Three distinct peaks were obtained for the substrate pNP-Xylose and both GS products, for which their respective peak areas were calculated. The peak area for pNP-Xylose in blank samples was used to normalize and estimate the concentrations of each product in the reaction samples.

**Electrospray Ionization Mass spectrometry (ESI-MS)**:

ESI-MS analysis was performed in a sensitivity mode on a high-resolution mass spectrometer (HRMS) containing quadrupole-TWIMS-TOF hybrid mass spectrometer (Synapt G2 HDMS; Waters Corp., Manchester, UK) in positive ionization mode with m/z scan range 100- 700 and from 300 -700.The applied experimental parameters were capillary voltage, 3.0 kV; sampling cone voltage, 40 V; source offset 80$^{o}$C ; source temperature, 100 °C; desolvation temperature 250$^{o}$C: Cone gas 50 L/ Hr and desolvation gas 600 L/Hr. Data analysis was performed using Mass Lynx 4.1

**Sample Introduction**:

GS reaction products were introduced directly into ESI-MS source based on the set parameters as mentioned above using an external manual syringe pump (Fisher Scientific) set at a flow rate of 600 µl/h. The spectra was collected for a period of 2 minutes. Based on the results shown in table 1 the molecular complex of the GS reaction products under different conditions produced an intense sodiated ion [M+Na]+with an m/z of 440.1135 which was consistently observed in all the reaction samples across the variants.Formation of cationic adducts with sugar molecules (in this case is sodium) are commonly formed in ESI-MS conditions when analyzed in positive mode due to the lack of protonated sites cationic adducts give stability to the sugar ions in gas phase and can further survive under ESI conditions. Mass spectrometry quickly confirmed the formation of the expected fucosylated disaccharide products, consistent with the designed glycosynthase reaction.

#### Liquid Chromatography - Mass Spectrometry (LC-MS)

Hydrophilic interaction liquid chromatography (HILIC) was performed on Acquity UPLC BEH-Amide,1.7-micron,2.1mmx100mm,(WatersCorporation,Milford,MA).Column temperature was set at 40$^{o}$C. The following mobile phases were used: A: 80:20 acetonitrile: water containing 0.1% Ammonium Hydroxide (LC-MS grade); B: 30:70 acetonitrile: water containing 0.1% Ammonium Hydroxide. The UPLC H-Class (Waters Corporation, Milford, MA) operated at a flow rate of 0.2 ml/min. 2µL of the sample was injected with an autosampler maintained at 10$^{o}$C. For the separation of compounds, the following gradient concentrations were used starting with 10% B for 2.0 minutes and later to 20% B for 4 min followed by 50 -70 %B for 10.0 minutes. Total run time of the LC-MS run was set for 12 minutes.

ESI-MS experiments were performed on high resolution mass spectrometer (HRMS) in a sensitivity mode using a quadrupole-TOF hybrid mass spectrometer (Xevo-G2- XS-Q-tof: Waters Corp., Manchester, UK) in negative ionization mode with m/z scan range 100–2000. The experimental parameters were set as follows: capillary voltage at 3.0 kV, sampling cone voltage at 30 V, source offset at 60°C, source temperature at 125°C, desolvation temperature at 250°C, cone gas flow rate at 50 L/hr, and desolvation gas flow rate at 600 L/hr. Data analysis was performed using MassLynx 4.1.

### Supplementary Results

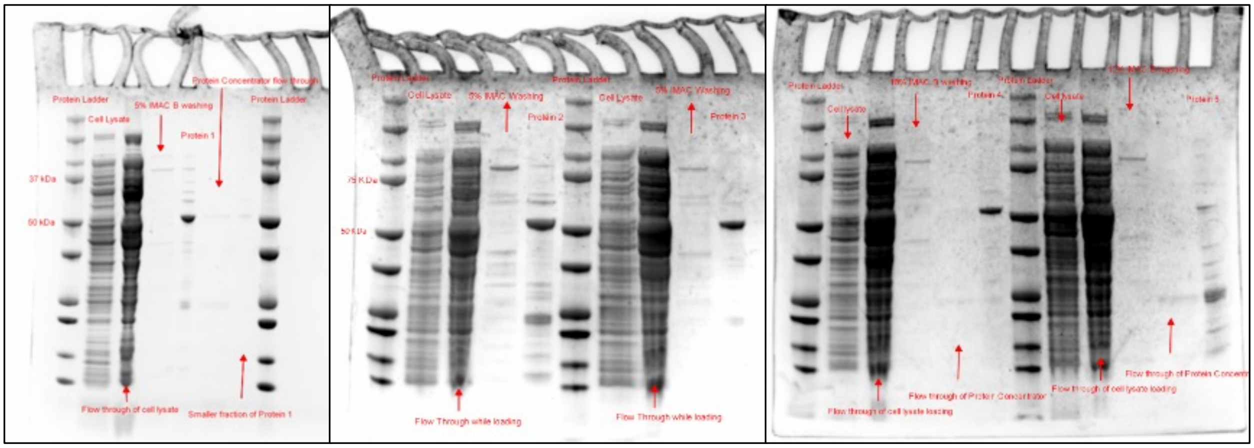


**Figure S1**. SDS–PAGE analysis of Ni-IMAC purification of GH29 α-fucosidase (*TmAfc*) variants expressed in *E. coli*; stain-free gels showing purification of the constructs, protein ladder, clarified cell lysate, flow-through during column loading; A dominant band at ~54 kDa corresponds to the His-tagged *TmAfc* and indicating effective capture and purification; faint additional bands reflect minor co-eluting host proteins.

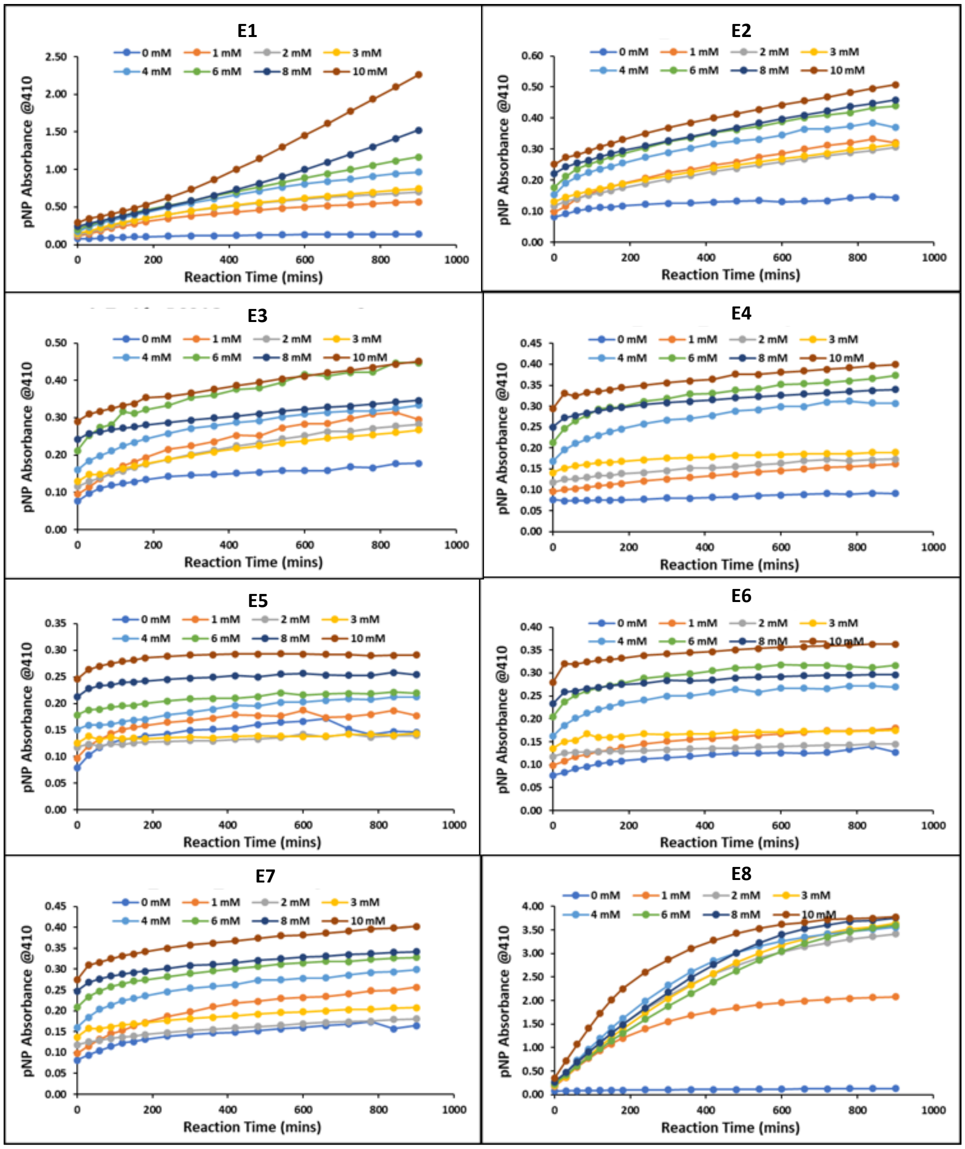


**Figure S2**. Progress curves for chemical-rescue assays of GH29 α-fucosidase variants. Time courses of p-nitrophenol release, monitored as $A_{410}$, at increasing substrate concentrations (0–10 mM pNP-α-L-fucopyranoside) for wild-type and representative nucleophile-deficient/engineered variants. Reactions were run under standard CR conditions (2 M $NaN_3$, 50 mM MES, pH 6.0; 60 °C), with absorbance recorded over the reaction time course. Higher substrate concentrations produce steeper initial slopes, reflecting increased initial rates. Initial velocities from the early linear regions were used for Michaelis–Menten analyses; 0 mM traces serve as baseline controls. Chemical rescue kinetics constants were determined by fitting the plots with the Michaelis-Menten equation. In Supplementary Figure S3 are shown example cases where kinetics parameters could be determined with certainty. For the rest of the purified proteins, it was not possible to determine the fitted kinetic parameters with certainty.

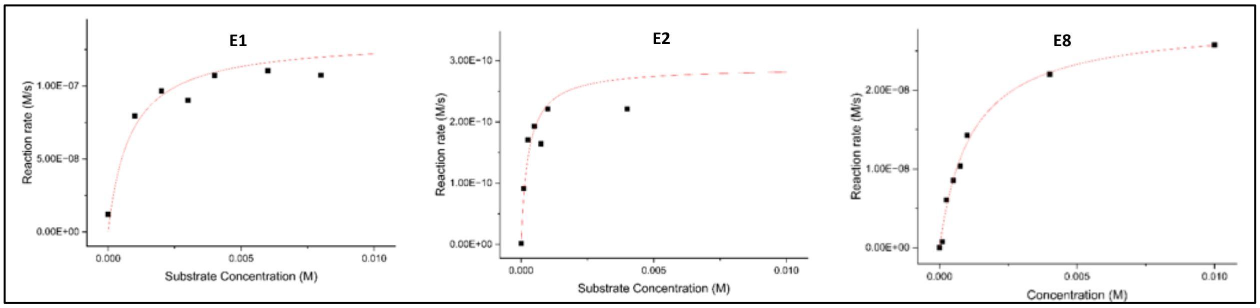


**Figure S3**. Michaelis–Menten kinetics for chemically rescued hydrolysis of GH29 α-fucosidase variants. Initial rates ($v_0$) were obtained from the early linear region of p-nitrophenol release ($A_{410}$) at increasing pNP-α-L-fucopyranoside concentrations (0–10 mM). Reactions were performed under standard CR conditions (50 mM MES, pH 6.0; 2 M NaN3; 60°C). Symbols denote measured $v_0$ values for wild-type E1, the D224G nucleophile-deficient parent E2, and representative engineered variants E8; dashed red curves are non-linear least-squares fits to the Michaelis–Menten model and goodness-of-fit ($R^2$) are summarized in the panel insets.

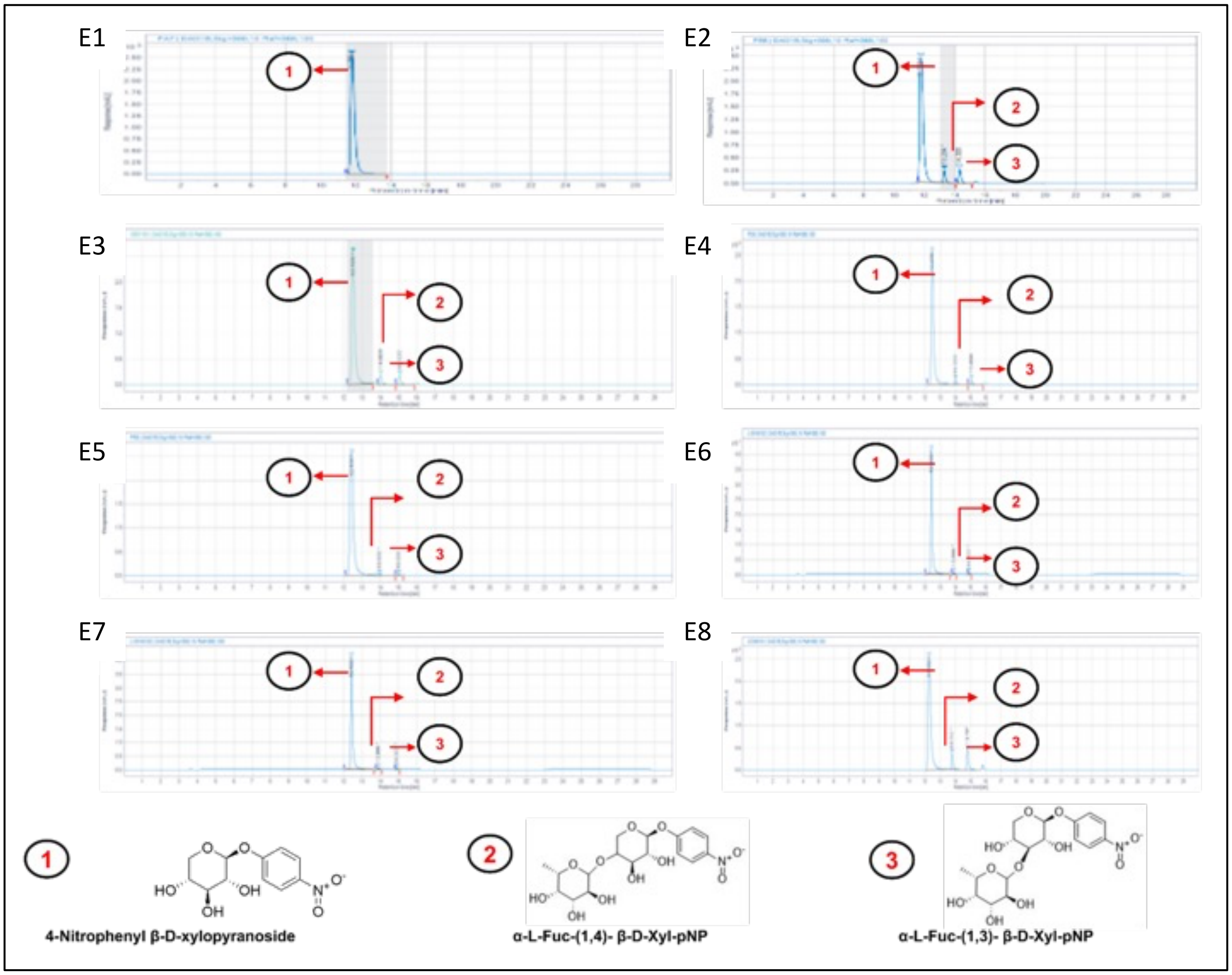


**Figure S4.** HPLC–UV analysis of glycosynthase reactions on pNP-β-D-xylopyranoside. Panels (a–h) show representative chromatograms for substrate-only control and selected purified enzyme variants of E1 to E8 after 24 h reactions. Peaks are annotated as: ① pNP-β-D-xylopyranoside (substrate), ② α-L-Fuc-(1→4)-β-D-Xyl-pNP, ③ α-L-Fuc-(1→3)-β-D-Xyl-pNP; detection at 254/300 nm, Product identities were confirmed with ESI-MS analyses.

| Mutants | Peak 1 (pNP-Xylose) | Peak 2 (1st GS Product) | Peak 3 (2nd GS Product) | Peak 2/Peak 3 Ratio |
|---|---|---|---|---|
| E1 | 100.00 | 0.00 | 0.00 | NA |
| E2 | 88.87 | 6.40 | 4.78 | 1.34 |
| E3 | 92.40 | 3.90 | 3.70 | 1.05 |
| E4 | 95.52 | 2.17 | 2.31 | 0.94 |
| E5 | 99.20 | 0.54 | 0.26 | 2.08 |
| E6 | 97.93 | 1.05 | 1.02 | 1.03 |
| E7 | 97.86 | 1.13 | 1.01 | 1.12 |
| E8 | 84.42 | 6.33 | 9.26 | 0.68 |

**Table T1**. HPLC–UV quantification of glycosynthase products across purified variants E1 to E8. Normalized peak areas (%) for Peak 1 = pNP-β-D-xylopyranoside (substrate), Peak 2 = α-L-Fuc-(1→4)-β-D-Xyl-pNP, and Peak 3 = α-L-Fuc-(1→3)-β-D-Xyl-pNP after 24 h reactions. Peak areas were integrated at 254/300 nm and normalized to the Peak 1 area measured in substrate-only blanks. The Peak 2/Peak 3 ratio reports relative regioselectivity; NA indicates no product detected above baseline.

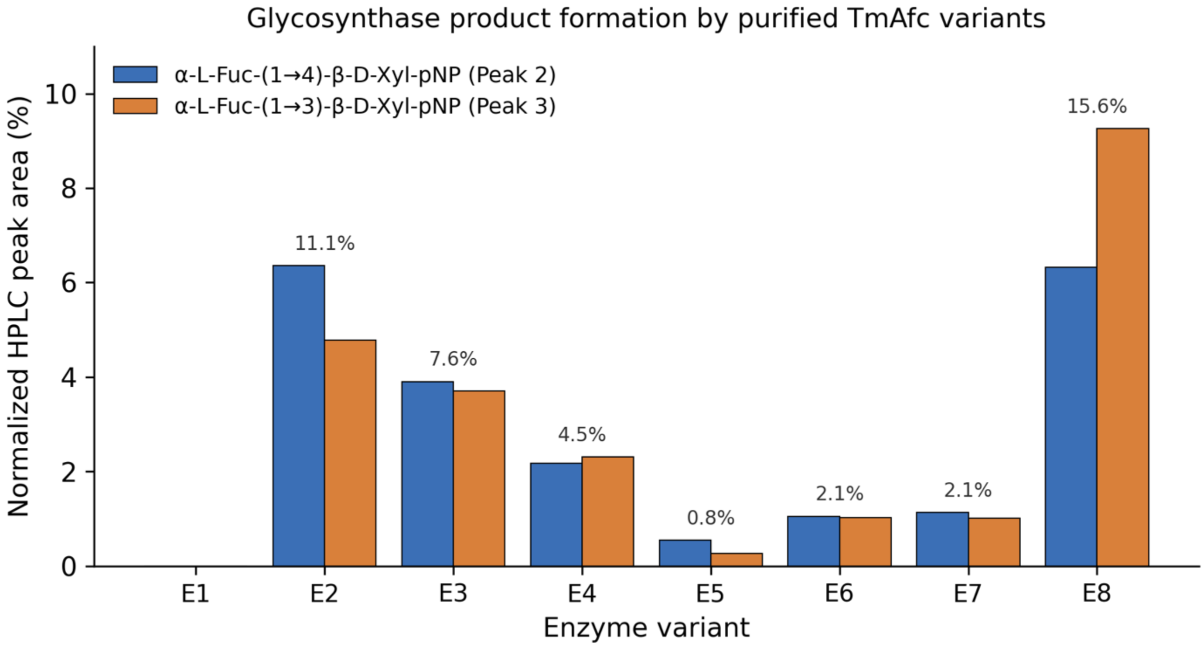


**Figure S5**. Regioisomer-resolved glycosynthase product yields for purified TmAfc variants. Normalized HPLC peak areas (%) are shown for the two fucosyl-xylose disaccharide products, α-L-Fuc-(1→4)-β-D-Xyl-pNP (blue) and α-L-Fuc-(1→3)-β-D-Xyl-pNP (orange), formed by the wild-type enzyme (E1) and engineered variants E2–E8 in glycosynthase reactions containing β-L-fucopyranosyl azide as the donor and pNP-xylose as the acceptor. The value above each group is the total product (sum of both regioisomers). No product was detected for the wild-type enzyme (E1), consistent with the absence of glycosynthase activity when the catalytic nucleophile is intact; E8 gave the highest total yield (15.6%), followed by the parent glycosynthase E2 (11.1%). Product regiochemistry was assigned following Cobucci et al. 2009.

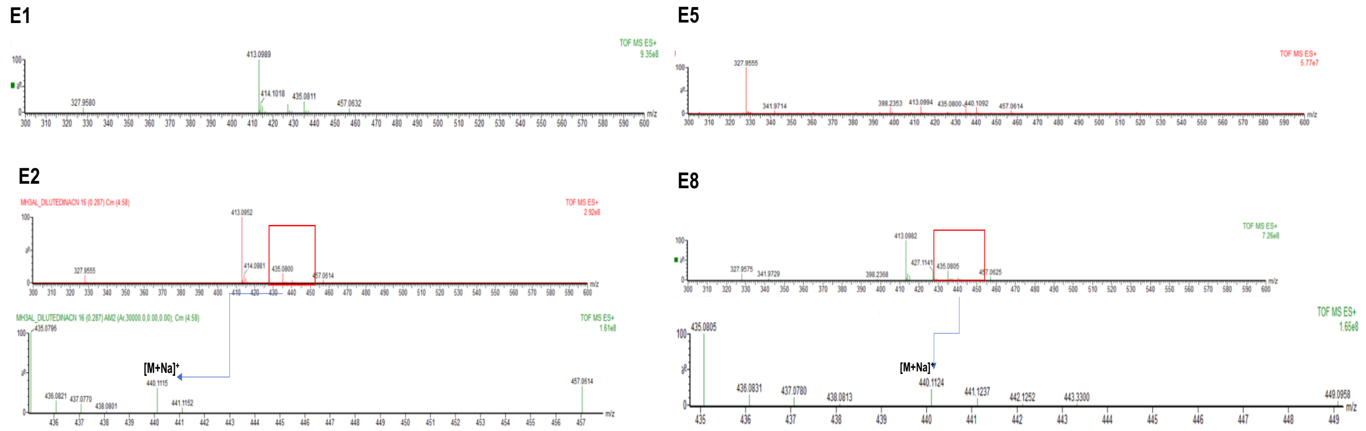


**Figure S6**. MS spectrum of glycosynthase reaction mixtures. Total ion current (TIC) traces and centroided mass spectra for a substrate-only control and representative enzyme variants (panel labels). The disaccharide product derived from fucosyl transfer to pNP-xylopyranoside is observed as the sodiated adduct [M+Na] ≈ m/z 440.116 (boxed) and is absent or near baseline in the control. Data were acquired on a Q-TOF in positive-ion mode (m/z 300–600) under standard source conditions; peak assignments are consistent with calculated exact masses and the HPLC–UV results (Fig.S4) ; E1 (wild type) and E5 mutant showed no presence of disaccharide product, whereas E2 and E8 enzyme showed presence of disaccharide product as expected.

| Samples | Intact Mol. Formula | Complex Observed | Observed mass. $[M+Na]^+$ | Theoretical mass $[M+Na]^+$ | Mass accuracy (ppm) |
|---|---|---|---|---|---|
| E1 | $C_{17}H_{23}NO_{11}$ | $C_{17}H_{23}NO_{11}Na^+$ | No detection (ND) | ND | ND |
| E2 | C17H$_{23}$NO11 | $C_{17}H_{23}NO_{11}Na^+$ | 440.1105 | 440.1163 | -13.0 |
| E3 | $C_{17}H_{23}NO_{11}$ | $C_{17}H_{23}NO_{11}Na^+$ | 440.1155 | 440.1163 | -1.81 |
| E4 | $C_{17}H_{23}NO_{11}$ | $C_{17}H_{23}NO_{11}Na^+$ | 440.1124 | 440.1163 | -8.8 |
| E5 | $C_{17}H_{23}NO_{11}$ | $C_{17}H_{23}NO_{11}Na^+$ | ND | 440.1163 | ND |
| E6 | $C_{17}H_{23}NO_{11}$ | $C_{17}H_{23}NO_{11}Na^+$ | ND | 440.1163 | ND |
| E7 | $C_{17}H_{23}NO_{11}$ | $C_{17}H_{23}NO_{11}Na^+$ | Very low | 440.1163 | ND |
| E8 | $C_{17}H_{23}NO_{11}$ | $C_{17}H_{23}NO_{11}Na^+$ | 440.1138 | 440.1163 | -5.6 |

**Table T2: Summary** of MS results of observed m/z, and calculated Δ ppm values of Glycosynthase reaction products: Based on the results shown in table 1. The molecular complex of the Glycosynthase reaction products under different conditions produced one intense sodiated ion [M+Na]+ m/z of 440.1135 which was consistently observed in all the reaction samples, with mass accuracy ranging between 1 and 13 ppm across variants. Formation of cationic adducts with sugar molecules (which in this case is sodium) generally seen in ESI-MS conditions when analyzed in positive mode.

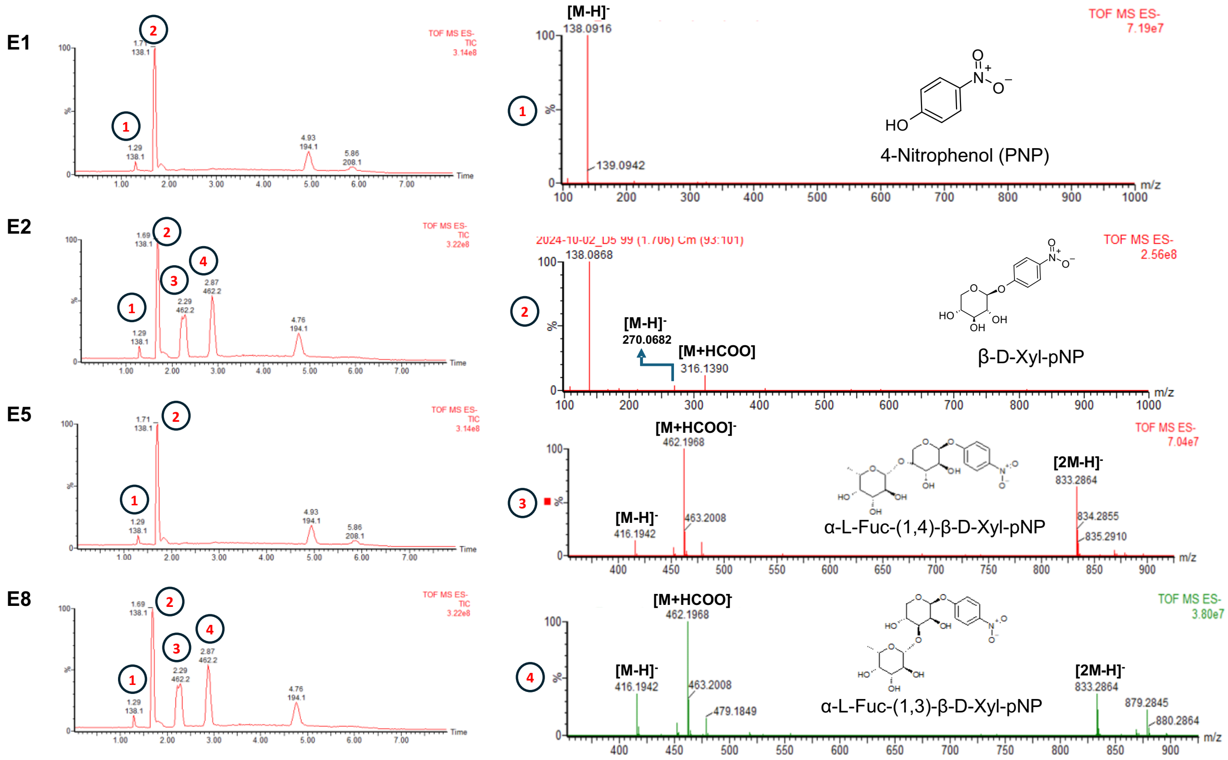

**Figure S7**: LC-MS analysis of glycosynthase (GS) reaction products from different mutants. Total ion chromatograms (TICs) for each mutant display distinct peaks corresponding to reaction products. **E1**.LC-MS shows two main peaks corresponding to deprotonated ion of 4-nitrophenol with an m/z 138 (1) and pNP-Fucose, m/z 284 (data not shown) **E2.** LC-MS presents additional peaks, with four distinct products : 4-nitrophenol (1), pNP-Fucose (2), α-L-Fuc-(1,4)-β-D-Xyl-pNP with m/z416 (3), and α-L-Fuc-(1,3)-β-D-Xyl-pNP with m/z 416 (4). **E5.** LC-MS primarily shows 4-nitrophenol (1) and pNP-Fucose (2) without further glycosylation products. **E8**. LC-MS also show four products, like E2. Structural representations of the identified products are shown alongside each panel, providing insight into the catalytic efficiency and specificity of each mutant.

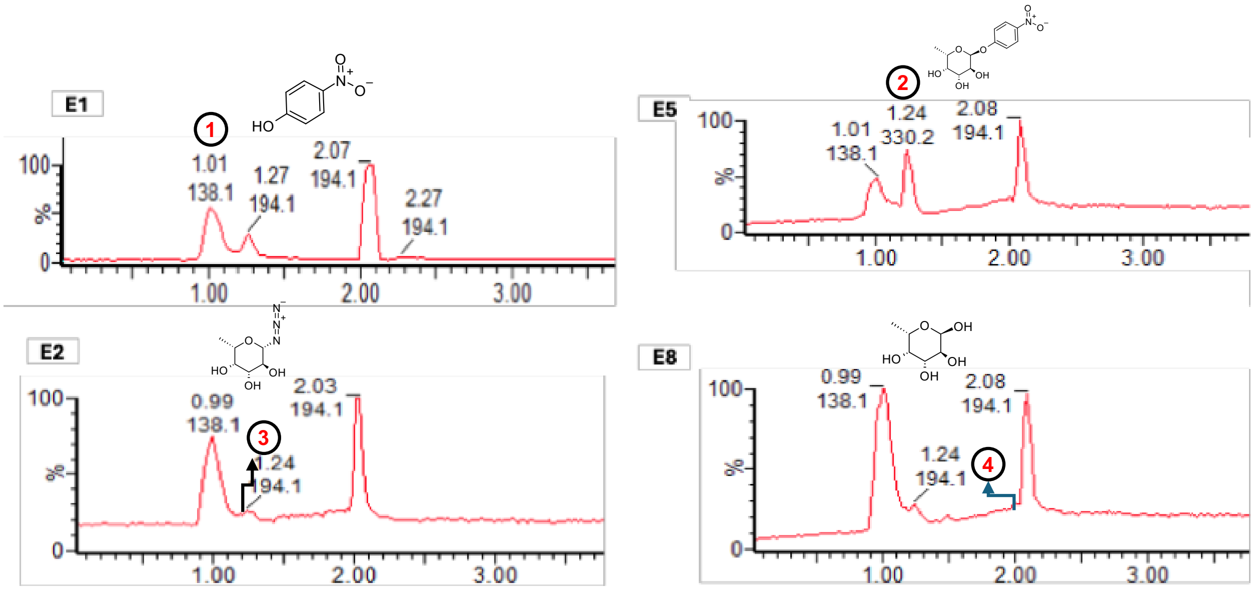


**Figure S8.** LC-MS analysis of chemical rescue (CR) reaction products across the enzyme panel. Total ion chromatograms (TICs; negative-ion mode) are shown for the wild-type enzyme (E1) and the engineered variants E2, E5, and E8, each incubated with pNP-fucose in the presence of 2 M sodium azide under standard CR conditions. Peaks were assigned by retention time (t_R) and accurate mass: Peak 1 (t_R ≈ 0.96-1.0 min), 4-nitrophenol (pNP) released from the substrate; Peak 2 (t_R ≈ 1.24 min), residual pNP-fucose; Peak 3 (t_R ≈ 1.46 min), β-L-fucopyranosyl azide (Fuc-N3); and Peak 4, free fucose arising from competing hydrolysis (t R ≈ 1.98 min). Fuc-$N_3$ is the activated glycosyl donor used in the glycosynthase reaction; its formation as a direct product of chemical rescue links the two reactions at the molecular level (Figure 4 ).

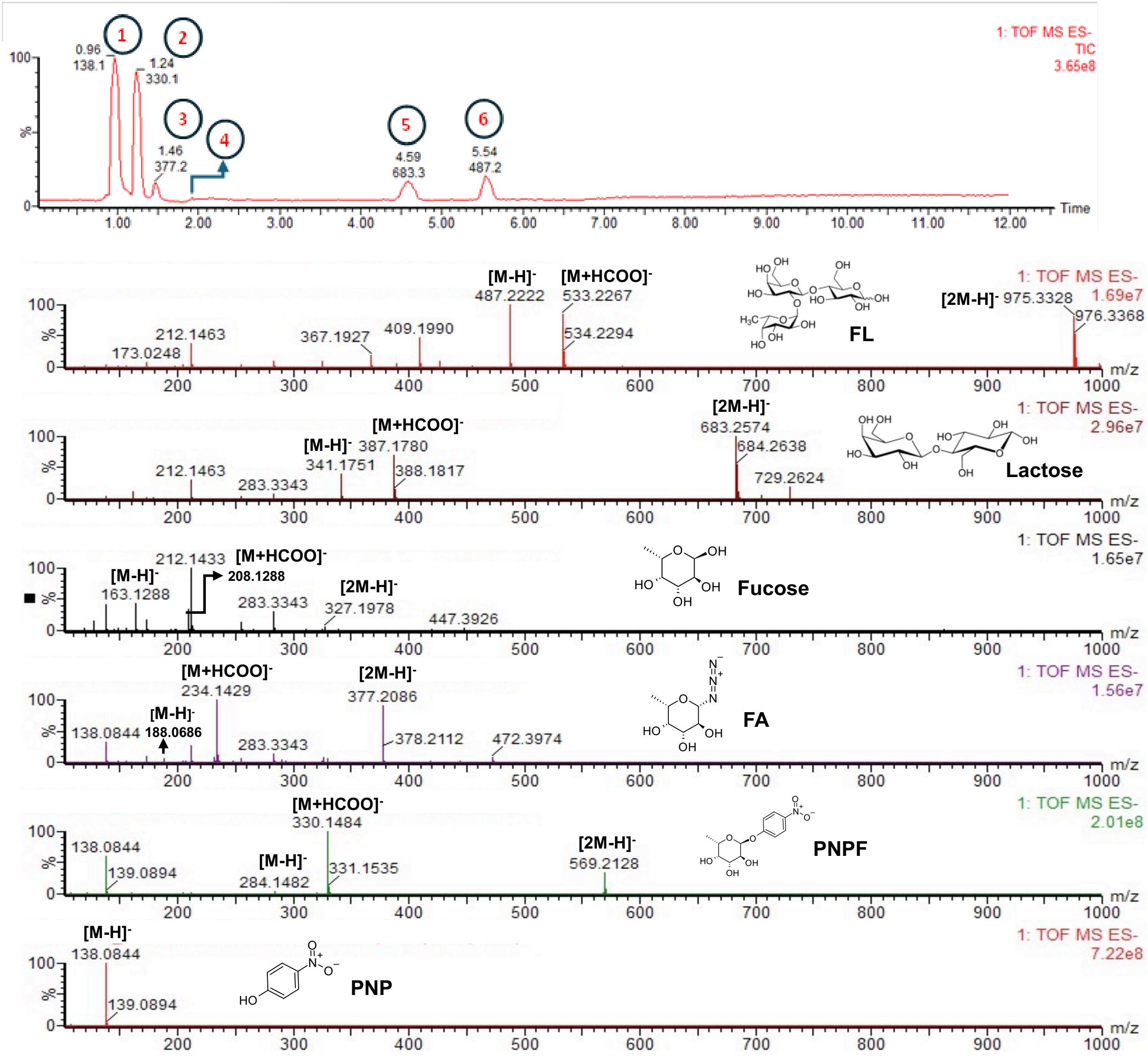


**Figure S9**. LC–MS profiling of standard solutions of sugars and nitrophenyl glycosides; UHPLC–QTOF total-ion chromatogram (TIC) with corresponding peaks 1–6 acquired in ESI-MS in negative mode with a scan range 100–1000 m/z. Annotations indicate the dominant ions for each peak (e.g.[M–H], [M+HCOO]$^-$ characteristic fragments and occasional deprotonated dimers; chemical structures next to spectra denote proposed assignments Peak 1. para-nitrophenol (PNP, mz/138), Peak 2. para- nitro phenol fucose (PNPF, m/z 284), Peak 3. Fucosyl azide (FA, m/z 188), Peak 4. Fucose (m/z 163), Peak 5. Lactose (m/z 341) and Peak 6. Fucosyl lactose (FL, m/z 487).LC separation and gradient conditions are as described in the LC–MS in the methods section. These data confirm compound identities and adduct/fragment patterns used for product assignment in the study and chemical rescue products were found at respective retention time and mass spectrum for all the enzymes as expected (Data not shown)

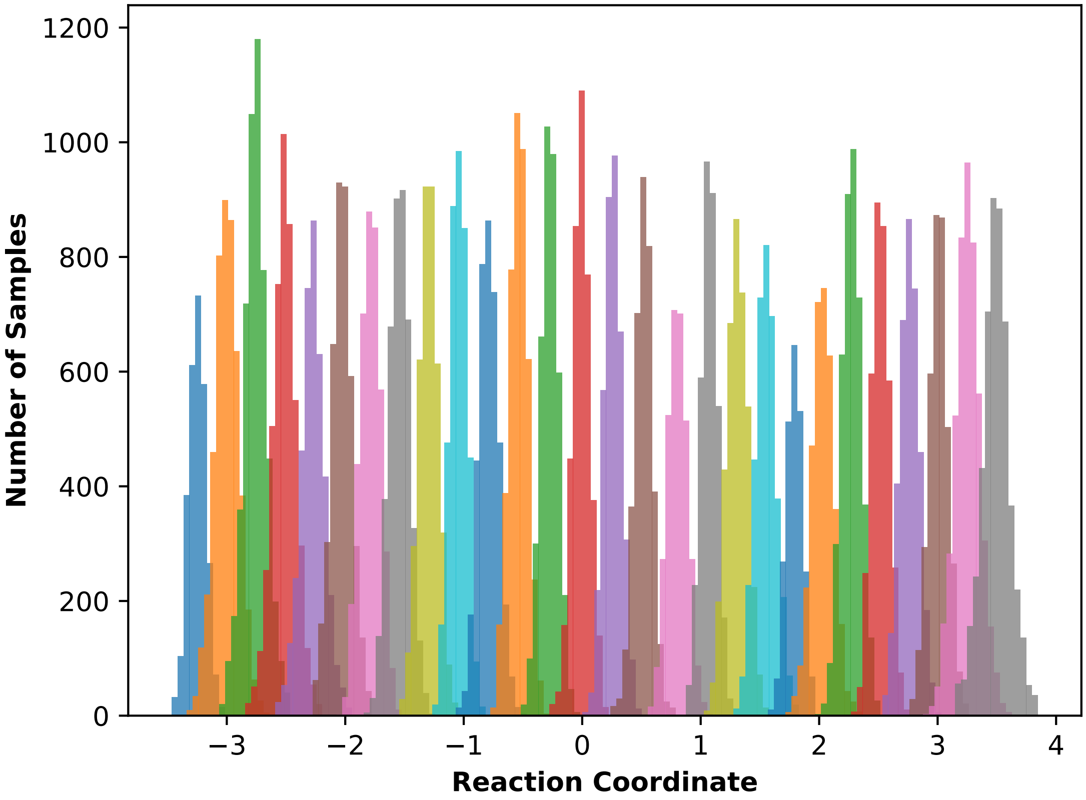


**Figure S10**: Umbrella-sampling window histograms for the QM/MM free-energy calculation of the chemical rescue reaction catalyzed by the TmAfc D224G glycosynthase. Each colored histogram shows the distribution of reaction-coordinate values sampled in one umbrella-sampling window during QM/MM (DFTB3) molecular dynamics of azide addition to pNP-fucose. The reaction coordinate is the likelihood-maximized linear combination of active-site collective variables identified by transition path sampling. Windows were spaced 0.25 units apart along the reaction coordinate and biased with a harmonic restraint (restraint weight 50 kcal/mol; five independent simulations per window). The substantial overlap between adjacent windows confirms continuous sampling across the entire reaction coordinate, allowing the histograms to be combined by the multistate Bennett acceptance ratio (MBAR) method into the continuous free-energy profile (potential of mean force) from which the activation free energy (ΔG‡ ≈ 8.7 kcal/mol) was obtained (see Figure 4j).
